# Aggregation recovers developmental plasticity in mouse polyploid embryos

**DOI:** 10.1101/263533

**Authors:** Hiroyuki Imai, Wataru Fujii, Ken Takeshi Kusakabe, Yasuo Kiso, Kiyoshi Kano

## Abstract

Polyploidy is comparatively prevalent in amphibians and fishes, but is infrequent in animals because of lethality after implantation. On the contrary, tetraploid embryos normally develop into blastocysts, and embryonic stem cells can be established from tetraploid blastocysts in mice. Thus, polyploidization does not seem to be so harmful during preimplantation development. However, the mechanisms by which early mammalian development accepts polyploidization are still poorly understood. In this study, we aimed to elucidate the effect of polyploidization on early mammalian development and to further comprehend its tolerability using hyperpolyploid embryos produced by artificial, repetitive whole genome duplication. Therefore, we successfully established several types of polyploid embryos (tetraploid, octaploid, and hexadecaploid), produced using repeated electrofusion of two-cell embryos in mice, and studied their developmental potential *in vitro*. We demonstrated that all types of these polyploid embryos maintained the ability to develop to the blastocyst stage, which implies that mammalian cells might have basic cellular functions in implanted embryos, despite polyploidization. However, the inner cell mass was absent in the hexadecaploid blastocysts. To complement the total cells in blastocysts, a fused hexadecaploid embryo was produced by aggregating a number of hexadecaploid embryos. The results indicated that the fused hexadecaploid embryo finally recovered pluripotent cells in blastocysts. Thus, our findings suggested that early mammalian embryos may have the tolerability and higher plasticity to adapt to hyperpolyploidization for blastocyst formation, despite intense alteration of the genome volume.

## INTRODUCTION

Polyploidy, produced by whole genome duplication, has some survival advantages, such as the increased ability to adapt to environmental changes, increased tolerance to genome damage, and increased flexibility and strength of tissues in plants and non-mammalian animals, which results in dramatic evolutionary changes, like the emergence of teleost fishes (Brodsky and Uryvaeva, 1985). Polyploidy is comparatively prevalent in amphibians and fishes; however, polyploid animals are infrequent because of lethality (Cleveland and Weber, 2014; Francesc et al., 2009; Lin et al., 1995). The mammalian genome has been suggested to show traces of two whole-genome duplication events (Kasahara, 2007; Ohno, 1970; Panopoulou and Poustka, 2005). In general, polyploidization is an infrequent phenomenon in present-day living mammals. Spontaneous duplication of the mammalian genome happens in less than 1% of fertilizations in mice, rats, rabbits, and pigs (Dyban and Baranov, 1987; McFeely, 1969). However, presently, polyploid mammals never develop to even newborn or adulthood, which indicates that present-day mammals do not have any plasticity for more polyploidization. In mice tetraploid embryos, the simplest whole genome duplication has been reported to develop to mid-gestation, but abort spontaneously at various developmental stages after implantation *in vivo* (Eakin and Behringer, 2003; James et al., 1992; Kaufman, 1991; Kaufman, 1992; Kaufman and Webb, 1990; Snow, 1975; Tarkowski et al., 1977), suggesting that a mechanism for discordance from alterations in the original genome volume exists during mammalian embryogenesis, especially post-implantation. However, the details of such a mechanism have not been fully elucidated. On the contrary, a preimplantation polyploid embryo has been suggested to have higher plasticity, even with various manipulations to them in mammals. With physical manipulation, a single blastomere, isolated from 2-, 4-, or 8-cell stage embryos, has the ability to develop to an entire blastocyst stage embryo, although of smaller size (Tarkowski, 1959; Tarkowski and Wroblewska, 1967). In terms of polyploidy, the plasticity seems to be maintained in early polyploid embryos, despite alteration of the genome volume. For example, although the number of total cells in mouse tetraploid blastocysts is lower than that in diploid blastocysts (Ishiguro et al., 2005; Kawaguchi et al., 2009), many studies have shown that tetraploid embryos can develop to the entire blastocyst stage embryo until implantation in standard embryo culture medium *in vitro* (Eakin and Behringer, 2003; Kawaguchi et al., 2009; Koizumi and Fukuta, 1995). In blastocysts, cell lineage into the embryonic or extra-embryonic tissue has already been programmed (Dietrich and Hiiragi, 2007). Inner cell mass has been detected in tetraploid blastocysts, and tetraploid embryonic stem cells have been successfully established from mouse tetraploid blastocysts (Imai et al., 2015). Tetraploid embryonic stem cells maintain intrinsic pluripotency and differentiation potential, including gene expression profiles and differentiation to germ layers, despite tetraploidization (Imai et al., 2015). The existence of a homeostatic mechanism controlling the cytoplasmic concentration of transcripts, in accordance with dramatic alterations in genome volume, caused by whole genome duplication, has also been suggested for tetraploid cells (Imai et al., 2016). These results indicate that whole genome duplication may not have an impact on the physiological progression of preimplantation embryo development in mammals. The purpose of this study was to examine whether hyperpolyploidy, produced by artificially repetitive whole genome duplication, affects early mammalian development, and to further comprehend the tolerability of polyploidization for sustaining plasticity for early mammalian development. In this study, we successfully produced a mouse full-term blastocyst with inner cell mass in a tetraploid, octaploid blastocyst. Even a hexadecaploid blastocyst embryo finally recovered pluripotent cells by aggregation of a number of embryos, suggesting that the intrinsic plasticity for blastocyst formation is basically maintained, despite intense alteration in the genome volume in mammals, even if the number of total cells can be compensated for in the preimplantation embryo.

## RESULTS AND DISCUSSION

### Establishment and characterization of hyperpolyploid embryos by repeated electrofusion

We first generated tetraploid mouse embryos by electrofusion of two-cell embryos of BDF1 male × BDF1 superovulated female matings and cultured these embryos to blastocyst embryos. To acquire basic knowledge about the effect of consecutive whole genome duplication on early mammalian development, we performed repeated cytoplasm electrofusion, as two-cell-stage embryos of tetraploids and octaploids were also electrofused to produce octaploids and hexadecaploids, respectively, to produce a hyperpolyploid embryo; these embryos were cultured for 4.5 days (Fig. 1A). Consequently, all of these produced polyploid embryos developed successfully into blastocyst stage embryos in a standard embryo culture medium *in vitro* by embryonic day 4.5 (Fig. 1B), which implies that mammalian polyploid cells may have normal cellular functions in early embryo formation before implantation. However, the blastocyst development rate was significantly lower in hexadecaploid embryos (56.1%), compared to those of the other polyploid embryos (diploid: 89.5%, tetraploid: 85.2%, and octaploid: 89.5%) (Table 1).

**Fig.1.**
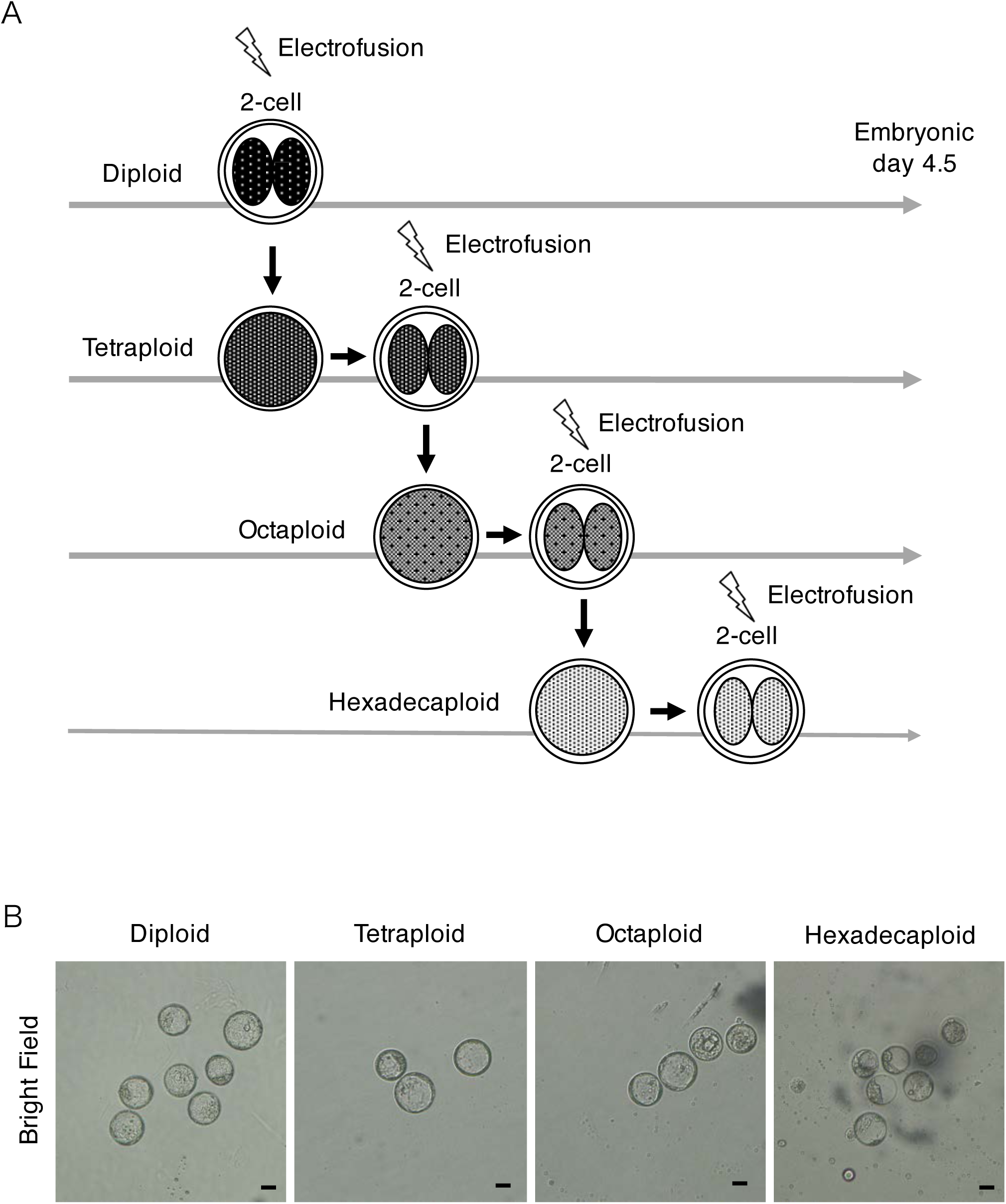
Hyperpolyploid mouse embryos. A) Generation scheme of mouse hyperpolyploid embryos by repeated electrofusion. The produced polyploid embryos were cultured for 4.5 embryonic days, and the analyses were performed. B) Morphology of mouse polyploid embryos at embryonic day 4.5. Each polyploid embryo developed into a blastocyst stage embryo with a blastocyst cavity. Scale bar; 50 μm.

**Table 1.** Developmental ratio of mouse polyploid embryos

|  | Normal (%) | Abnormal (%) | total |
| --- | --- | --- | --- |
| Diploid | 77 (89.5) | 9 (10.5) | 86 |
| Tetraploid | 52 (85.2) | 9 (14.8) | 61 |
| Octaploid | 59 (89.5) | 10 (14.5) | 69 |
| Hexadecaploid | 32 (56.1*) | 25 (43.9) | 57 |

\*: $p < 0.01$

Tetraploid mouse embryos, the simplest whole genome duplication embryo, rarely occur in nature, but they can be classically produced by methods that chemically block cell division, such as through the use of cytochalasin B (Snow, 1973). Recently, electrofusion of two-cell-stage embryos by electrical stimulation has been popular and convenient for efficient production of polyploid embryos in mice (Berg, 1982; Kubiak and Tarkowski, 1985; Kurischko and Berg, 1986). We succeeded in establishing a mouse tetraploid, octaploid, and even a hexadecaploid embryo, by consecutive electrofusion of the two-cell stage embryos of diploids, tetraploids, and octaploids, respectively. In particular, a hexadecaploid embryo was successfully established in mice for the first time. These results suggested that although the genome volume of the hexadecaploid cell was eight-fold, compared to that of a diploid cell, hyperpolyploid embryos, such as the hexadecaploid embryo, still maintain the ability to develop to blastocyst stage embryos similar to other lesser polyploid embryos. Consistent with these results, our experiments also showed that the developmental ratio decreased significantly in the hexadecaploid embryos. The development of octaploid embryos is affected by an increase in apoptosis, autophagy, and epigenetic modification (Wu et al., 2017), which indicates that higher the ploidy level, more severe the damage to embryogenesis from these intracellular alterations in the early development of a hexadecaploid embryo, compared to diploid and lesser ploidy embryos.

### Absence of inner cell mass in the hexadecaploid embryo

To acquire molecular information regarding key factors of the blastocyst, indirect immunofluorescence and gene expression analyses were performed in these polyploid blastocysts (Fig. 2A, B, and C). Indirect immunofluorescence for CDX2 (a trophectoderm marker) of blastocyst stage embryos (embryonic day 4.5) indicated that trophectoderm-specific molecules were expressed at the protein level in all polyploid blastocysts, similar to their expression in diploids (Fig. 2A). We also performed indirect immunofluorescence for OCT3/4 (an inner cell mass marker), which localizes in tetraploid and octaploid blastocysts, but not in hexadecaploid blastocysts (Fig. 2B). To elucidate the transcriptional levels of these various polyploidy levels in the blastocyst embryo in detail, we performed RT-PCR analyses for *Gata6* (a primitive endoderm marker) and *Nanog* (an epiblast marker), in addition to *Cdx2* and *Oct3/4*. The transcriptional analyses indicated that *Cdx2* mRNA was detected in all polyploid embryos; however, the inner cell marker genes, *Oct3/4*, *Nanog*, and *Gata6*, were not or were only slightly expressed in the hexadecaploid blastocysts (Fig. 2C) The data implies that the inner cell mass was absent in hexadecaploid blastocysts in mice.

**Fig.2.**
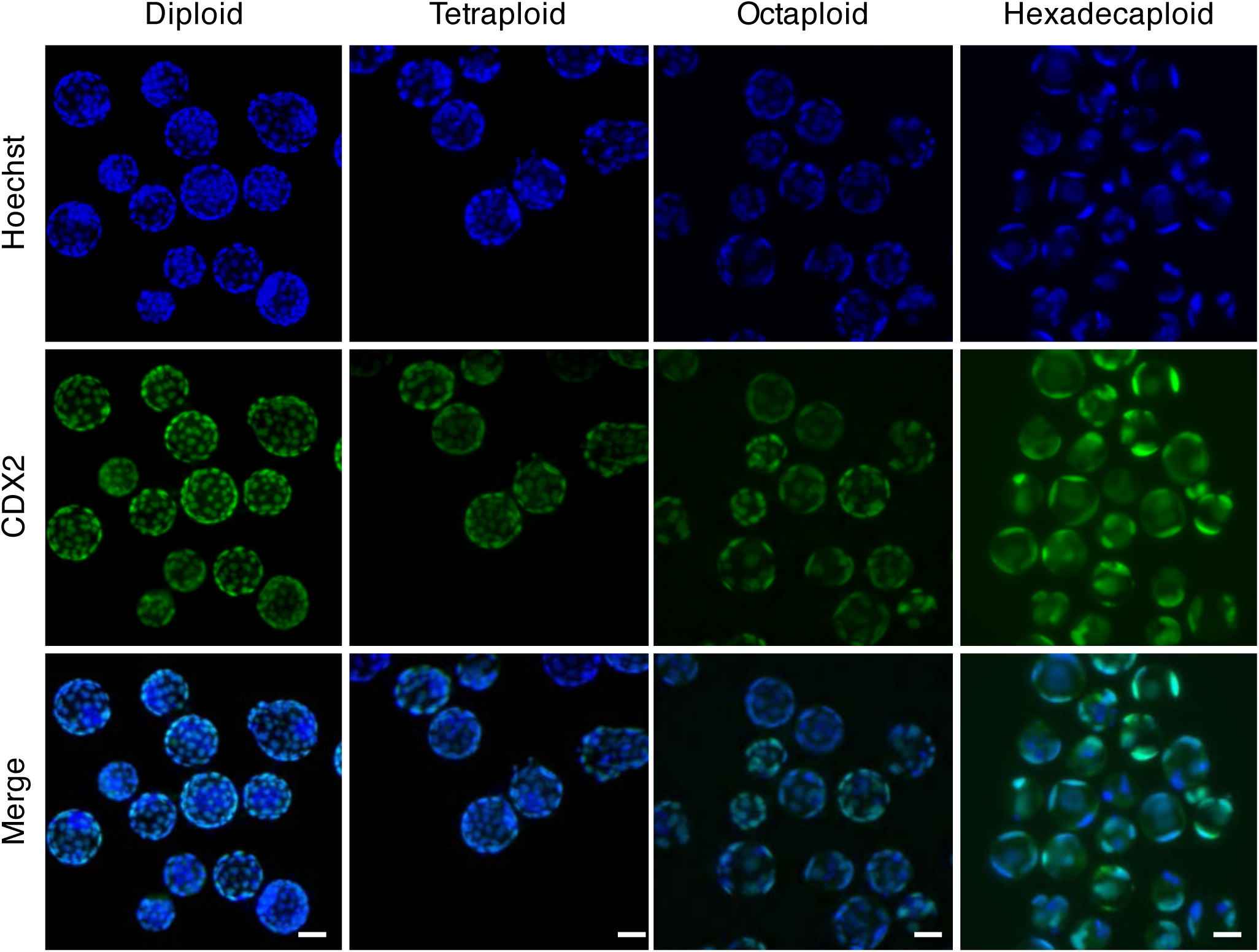
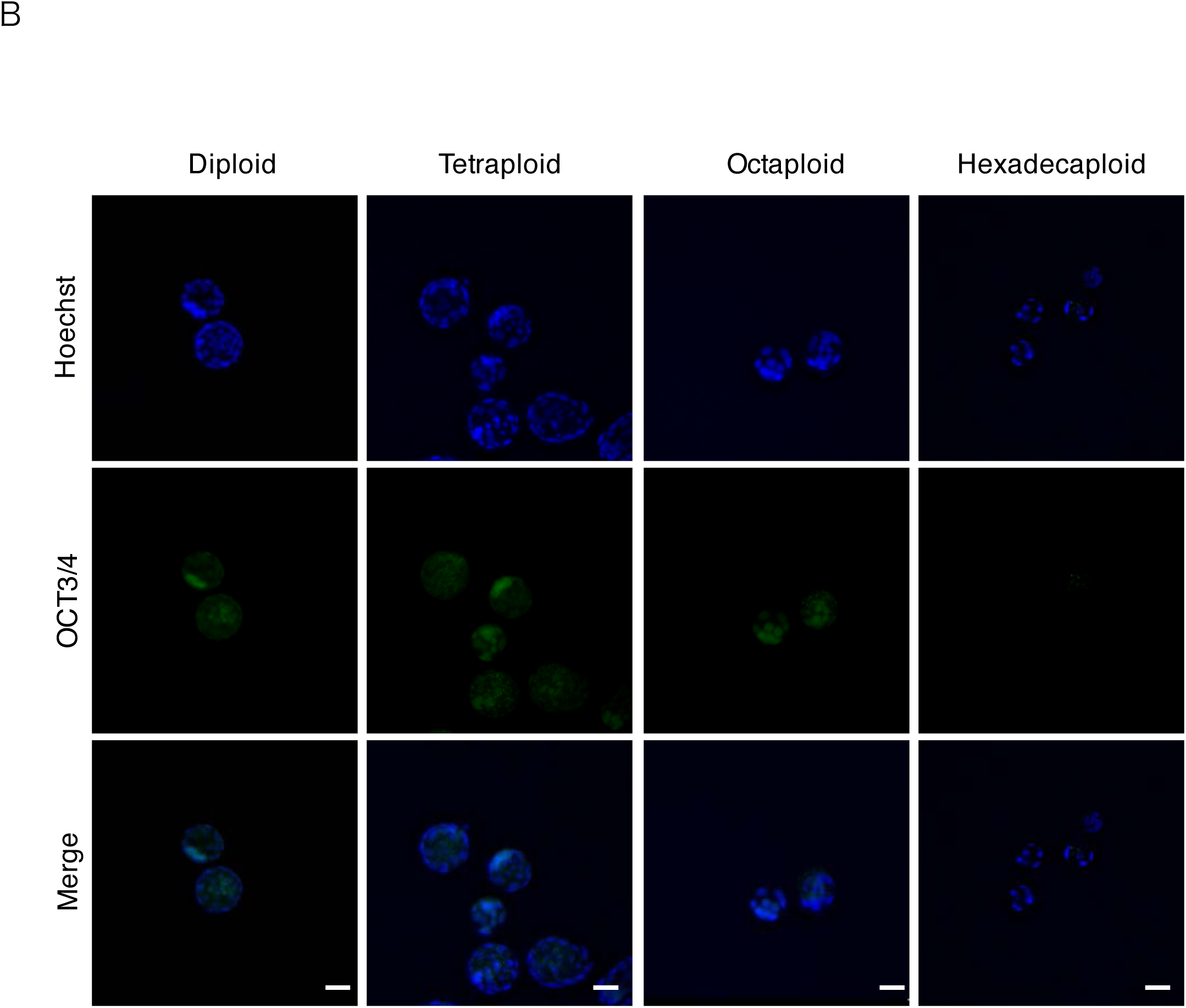
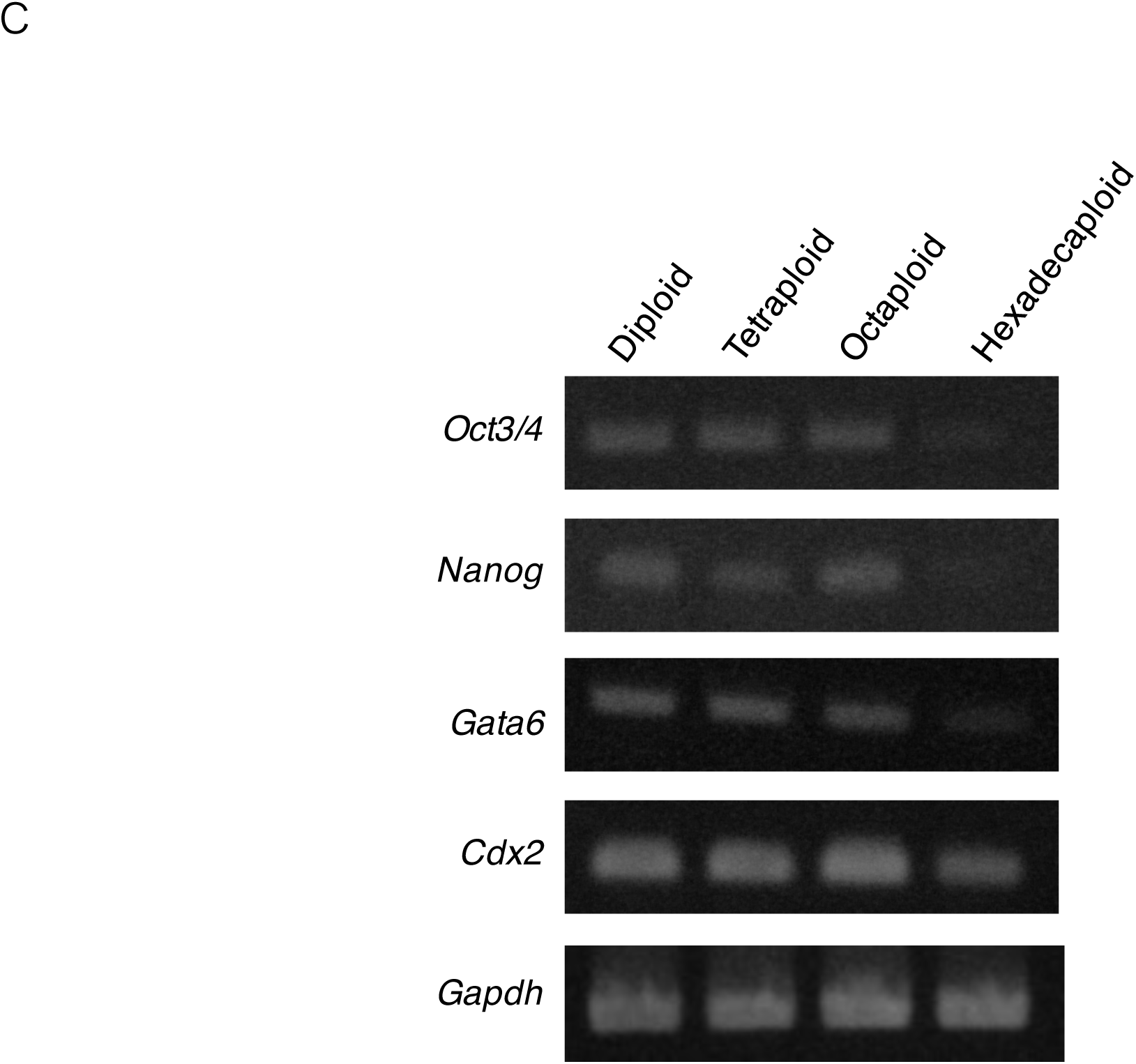
Localization and expression analyses of molecular markers of blastocysts in mouse polyploid embryos at embryonic day 4.5. A) Immunostaining for CDX2 (TE; Trophectoderm) and OCT3/4 (ICM; Inner Cell Mass) in mouse polyploid embryos. CDX2 was detected in all polyploid embryos. OCT3/4 was not detected in the hexadecaploid embryos. DNA was stained with Hoechst 33342. Scale bar; 50 μm. B) RT-PCR analysis of the key factors of blastocyst genes: *Oct3/4*, *Nanog*, and *Gata6* for inner cell mass, *Cdx2* for the trophectoderm, and *Gapdh* as a control in various polyploid embryos. *Oct3/4*, *Nanog*, and *Gata6* were not expressed in the hexadecaploid embryos.

Tetraploid blastocysts can be classified into two groups, based on the presence or absence of the inner cell mass in mice (Wen et al., 2014). If the number of cells in an embryo is artificially increased in the four-cell stage embryo, the ratio for the presence of the inner cell mass increases in tetraploid embryos (Wen et al., 2014). In mouse octaploid embryos, the number of cells with inner cell mass decreases significantly, and these cells also have more variety compared to that of diploid embryos in blastocysts (Wu et al., 2017). We have successfully established tetraploid embryonic stem cells (TESCs) from mouse tetraploid blastocysts; however, the efficiency of the establishment of TESCs (15%; Table 2) was quite low compared to that of ESCs (88 %; Table 2) (Imai et al., 2015). We have shown that the substantial tolerability for the maintenance of the characterization as embryonic stem cells is sustained, despite an increase in the genome volume in mouse tetraploid embryonic stem cells (Imai et al., 2015).

**Table 2.**
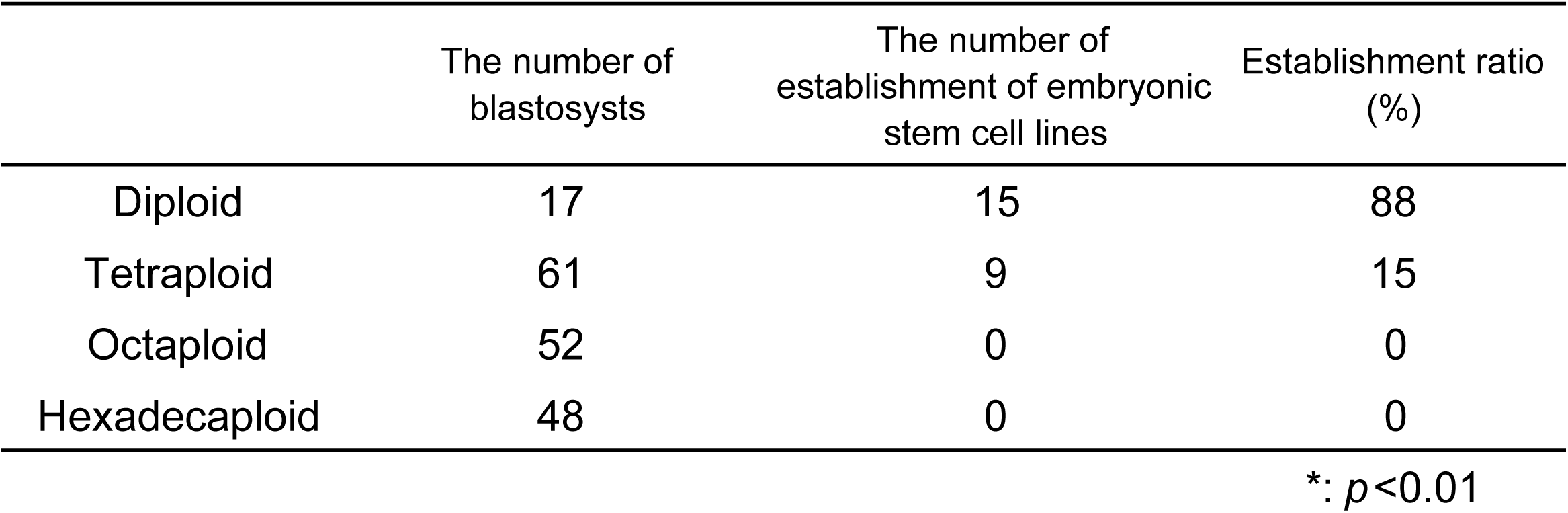
Establishment ratio of mouse polyploid embryonic stem cells

Embryonic stem cells established from the inner cell mass can sustain the pluripotency for differentiation, and can be induced to differentiate into all types of tissues, including germ cells, under specific culture conditions (Evans and Kaufman, 1981; Hayashi et al., 2012; Hayashi et al., 2011). We tried to establish embryonic stem cells from mouse octaploid and hexadecaploid blastocysts; however, we were never successful in establishing mouse octaploid and hexadecaploid embryonic stem cells from octaploid and hexadecaploid blastocysts, respectively (Table 2). These studies imply that some tetraploid blastocysts only possess enough cells in the inner cell mass for establishment of embryonic stem cells; however, octaploid and hexadecaploid blastocysts might be crucially insufficient or absent for the establishment of their embryonic stem cells. Our data showed that the inner cell mass was absent in both the proteins and transcripts in hyperploid, hexadecaploid blastocysts, which indicates two possibilities. The first is a massive insufficiency in the total cell numbers, because the cell number of hexadecaploid blastocysts is approximately eight parts of that of diploid blastocysts. The other possibility for the disappearance of the inner cell mass in hexadecaploid blastocysts is that the larger genome volume might impair intrinsic acquirement of pluripotency through the inner cell mass.

### Aggregation recovers the inner cell mass in hexadecaploid embryo

To test these hypotheses for the absence of the inner cell mass in hexadecaploid embryos, we first produced an aggregated embryo from hexadecaploid embryos. Four separated hexadecaploid embryos, whose zona pellucida was removed, were aggregated together in a small hole, made by a needle on a plastic culture dish for physical attachment, and the aggregated hexadecaploid embryo was cultured until embryonic day 4.5 (Fig. 3A). After 12 hours of culture, the resulting aggregated hexadecaploid embryos continued to develop into a single morula embryo (Fig. 3B). Indirect immunofluorescence indicated that for the localization of OCT3/4, inner cell mass-specific molecules were recovered at the protein level in the aggregated hexadecaploid blastocysts at the inner cell mass, similar to the localization of these in diploid, tetraploid, and octaploid embryos (Fig. 3C). We further confirmed the transcriptional levels of inner cell mass-specific molecules in the aggregated hexadecaploid blastocysts using RT-PCR analysis for the *Oct3/4, Nanog, Gata6*, and *Cdx2*, which indicated that the expression levels of the *Oct3/4*, *Nanog,*and *Gata6* were clearly recovered in the aggregated hexadecaploid blastocysts, such as in the other lesser polyploid blastocysts, compared to the non-aggregated hexadecaploid blastocysts (Fig. 3D). These results indicated that the inner cell mass might be restored in blastocyst-stage embryos by compensating for the number of total cells in the multiple hexadecaploid aggregation.

**Fig.3.**
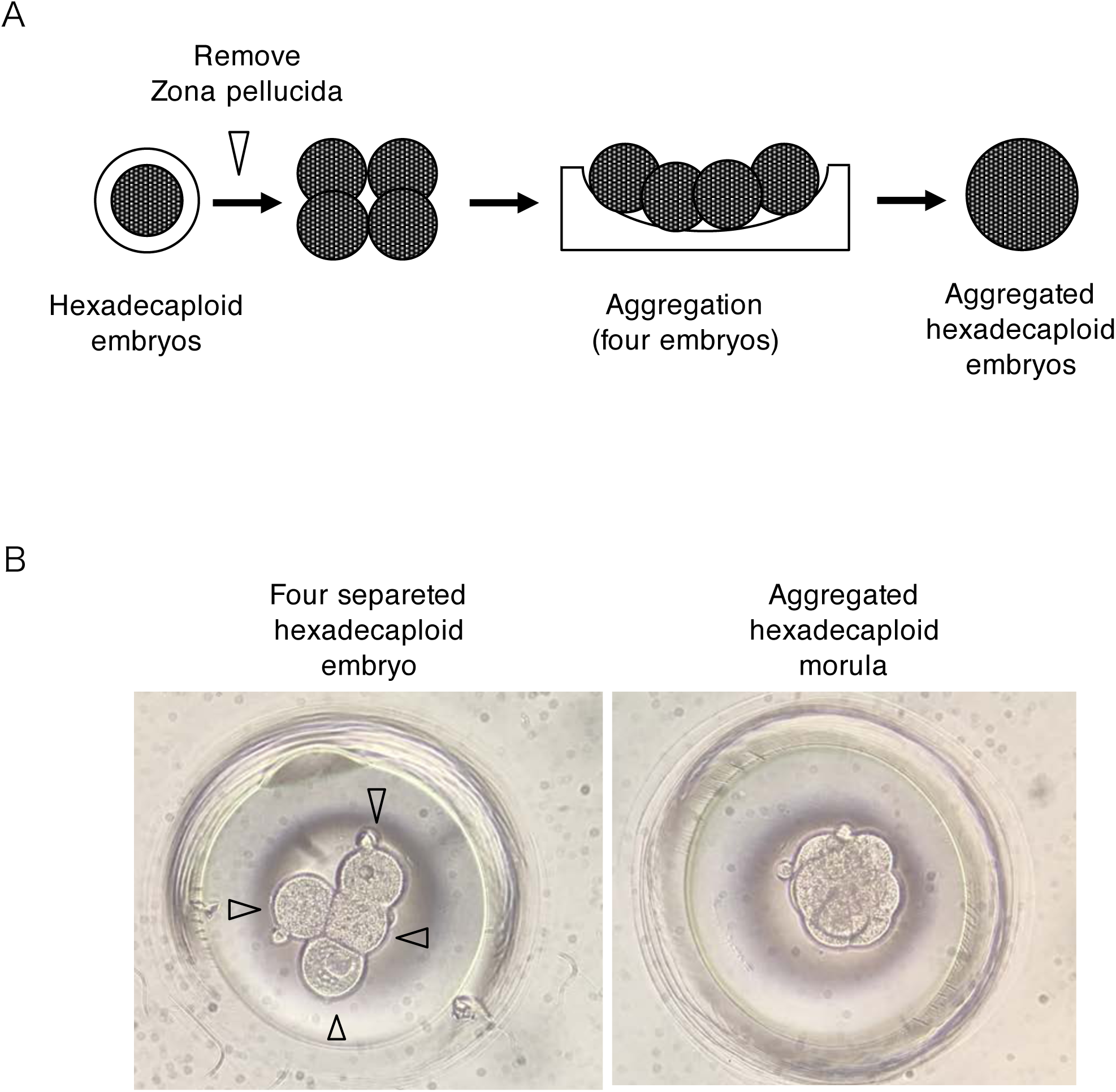
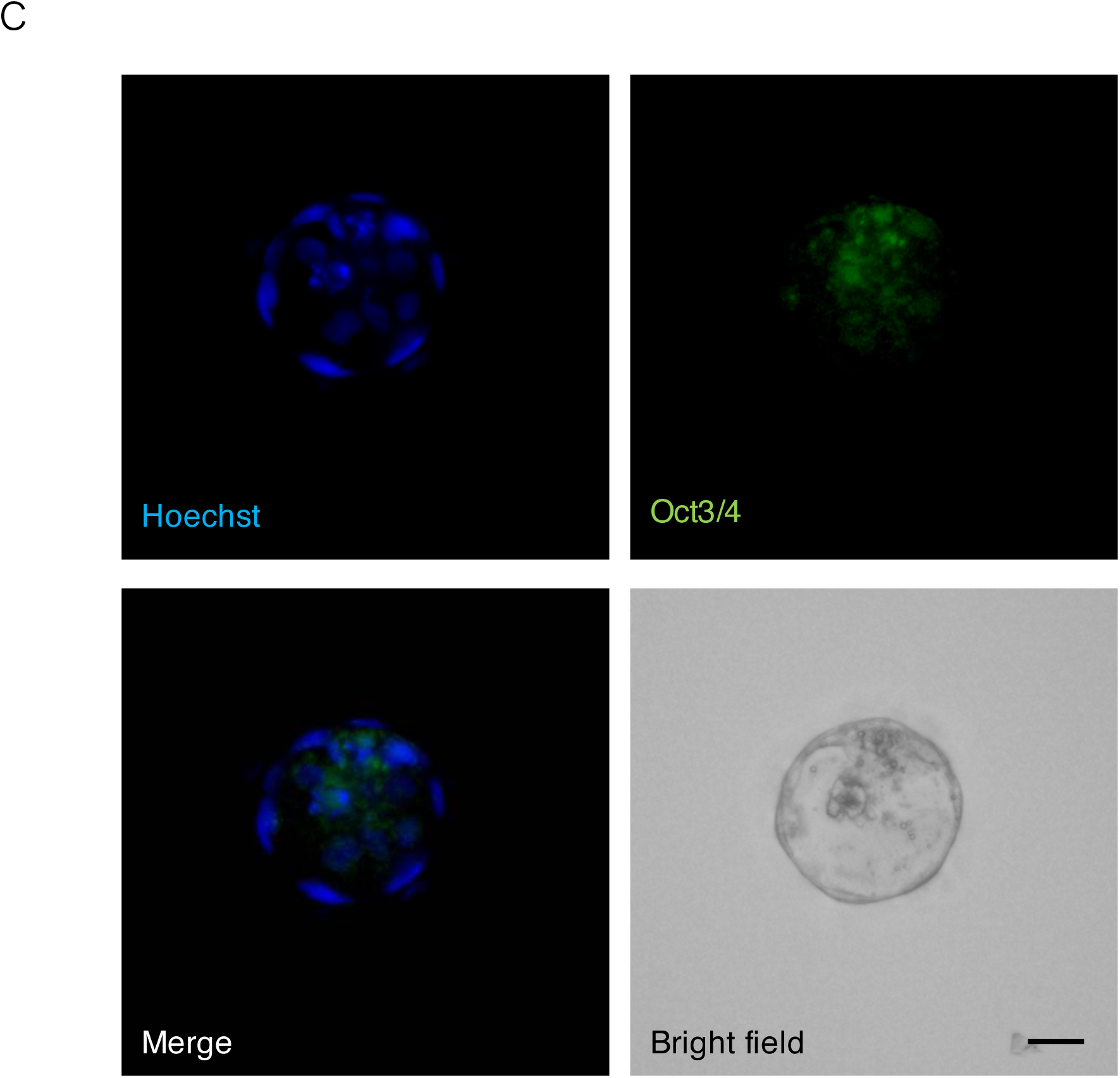
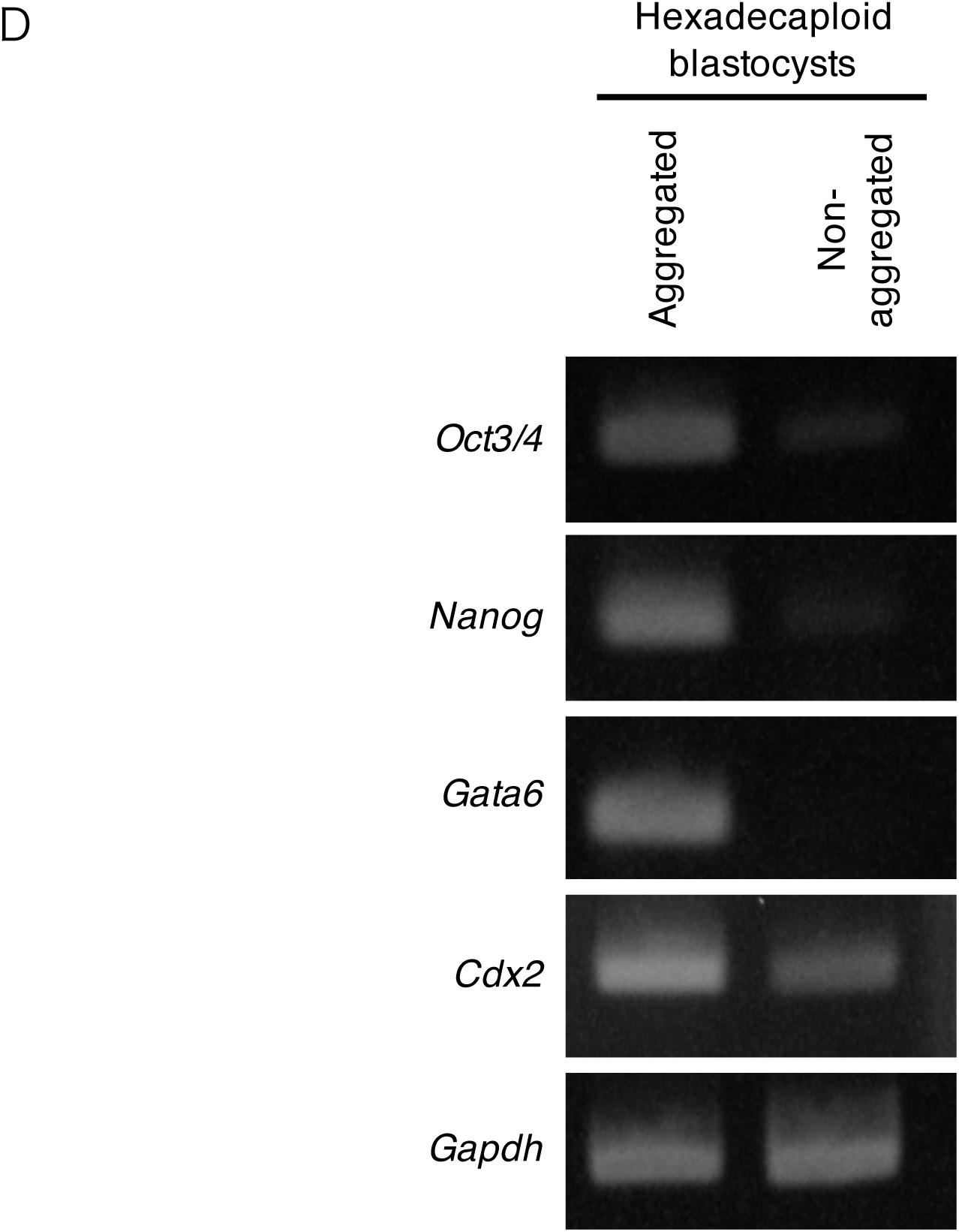
Compensation for the number of total cells for development in mouse hexadecaploid embryos. A) Method scheme for generation of multiple aggregated hexadecaploid embryos. After removing the zona pellucida, four separated hexadecaploid embryos were aggregated in a hole and cultured until embryonic day 4.5. B) Bright field images of the aggregated hexadecaploid embryo. Four separated hexadecaploid embryos (left; arrow head) were cultured in a small hole. After 12 hours, these hexadecaploid embryos were fused as a single embryo, and the aggregated embryo finally developed into a morula embryo (right). C) Immunofluorescence of OCT3/4 in the aggregated hexadecaploid embryo. OCT3/4 was localized at the inner cell mass of the aggregated hexadecaploid blastocysts. DNA was stained with Hoechst 33342. Scale bar; 50 μm. DRT-PCR analyses of the aggregated hexadecaploid blastocyst. *Oct3/4*, *Nanog*, and *Gata6* genes were clearly recovered in the aggregated hexadecaploid blastocysts, compared to the non-aggregated hexadecaploid blastocysts.

Mouse tetraploid embryos can develop normally until implantation and retain properties for the expression of pluripotent gene markers [our unpublished data, (Eakin and Behringer, 2003; Kawaguchi et al., 2009; Koizumi and Fukuta, 1995; Pfeiffer et al., 2013; Pralong et al., 2005). The number of total cells in the tetraploid blastocysts is lower, around one-half, compared to those in diploid blastocysts (Ishiguro et al., 2005; Kawaguchi et al., 2009). The two distinct cell lineages, the inner cell mass or the trophectoderm, are differentiated in the mammalian blastocyst. Three controversial models—pre-patterning, inside-outside, and cell polarity—have been proposed for the differentiation of the two distinct cell lineages during pre-implantation development (Wennekamp et al., 2013). The inside-outside model (Tarkowski and Wroblewska, 1967) proposes that the position of a cell, rather than any cell-intrinsic differences, induces the cell fate of the morula-stage embryo, where cells on the inside form the inner cell mass, and cells on the outside form a trophectoderm. Among these models, the inside-outside model could be more suitable for explaining why the inner cell mass emerges in tetraploid and octaploid blastocysts, but not in the hexadecaploid blastocysts. In general, the higher the ploidy, the larger the cell size and the lower the total cell number in early stage embryos. We speculated that the number of total cells composing an embryo is just sufficient to form the inside and outside cells in tetraploid and octaploid morula, but extremely insufficient in hyper-ploidy embryos, such as the hexadecaploid morula, which consists of only a few cells. Our study on the aggregated hexadecaploid embryos demonstrated that restitution of the number of total cells could be quite important for embryo differentiation into the two cell lineages; the trophectoderm and especially the inner cell mass. Buemo *et al*. also revealed that embryo aggregation in pigs improves the cloning efficiency and embryo quality, which also indicates that the number of total cells is a factor for the early mammalian embryo (Buemo et al., 2016).

Another possibility, in which hyperploid cells might impair the intrinsic acquirement of pluripotency as inner cell mass, has been continuously argued. However, further studies are needed to clarify the characteristics of intrinsic pluripotency of hyperploid cells in mammals, because we failed to establish embryonic stem cells from oxtaploid and hexadecaploid blastocysts. In the future, experiments for the establishment of embryonic stem cells from aggregated hyperploid blastocysts could clarify that mammalian cells have unexpected plasticity for early embryo development in mammals, despite hyper-duplication of the genome volume of a single cell.

## Conclusions

In this study, we produced hyperpolyploid embryos by consecutive electrofusion in mice to examine whether hyperpolyploidization affects early mammalian development and to further comprehend the tolerability of polyploidization in sustaining plasticity for early mammalian development. We found that the inner cell mass was absent in a single hexadecaploid blastocyst; however, complementing the number of total cells of the embryo made the early mouse embryo recover pluripotent cells in the hexadecaploid blastocyst. These results indicated that a preimplantation embryo may have a higher plasticity to adapt to repeated polyploidization, if the number of total cells of an embryo is enough to organize the whole blastocyst embryo in the preimplantation stage of mammals. Our data also implies that the early mammalian embryo may have the tolerability and higher plasticity to adapt to hyperpolyploidization, if conditions such as the number of total cells of an embryo are sufficiently qualified for the maintenance of blastocyst differentiation.

## MATERIALS AND METHODS

### Production of polyploid embryos

All mice used in this study were bred at Yamaguchi University. All procedures were approved by the Yamaguchi University IACUC. BDF1 female mice were superovulated for the generation of tetraploid embryos. Following human chorionic gonadotropin injection, the females were mated with BDF1 males. Two-cell-stage embryos were collected and electrofused as previously described (Imai et al., 2015). To produce octaploid embryos, fused two-cell-stage tetraploid embryos were selected and electrofused again (Bex). To produce hexadecaploid embryos, the same procedure was conducted. To produce an aggregated hexadecaploid embryo, after removal of the zona pellucida with acidic Tyrode’s solution (Sigma), four hexadecaploids were placed on and adhered to a hole made by an aggregation needle (BLS) on a culture dish.

### Immunocytochemistry

At embryonic day 4.5, the embryos were fixed with 0.5% Polyvinylpyrrolidone and 4% paraformaldehyde in PBST. After permeabilization with 0.5% PVP and 0.5% Triton-X 100 in PBS for 30 min and blocking with 0.5% PVP and 10% serum in PBS for 30 min, the embryos were incubated with the primary antibodies shown in Table S1 at 4°C overnight and then with secondary antibodies at 25°C for 60 min. After washing with PBS, the embryos were stained with 5 μg/mL Hoechst 33342 in PBS and then observed using a confocal scanning microscope (Keyence).

### Reverse transcription polymerase chain reaction

For reverse transcription polymerase chain reaction (RT-PCR), total RNA was isolated from embryos using the ReliaPrep RNA Cell MiniPrep System (Promega). The total RNA of each sample was quantitatively measured using a spectrophotometer (Thermo Fisher Scientific), and the cDNA was synthesized from 1 μg of total RNA with a QuantiTect Reverse Transcription Kit (Qiagen). Table S2 lists the primers used to detect *Gapdh*, *Nanog*, *Oct3/4*, *Cdx2*, and *Gata6*. RT-PCR was performed using a PrimeSTAR HS (Takara) with a 20 μL volume and 1 μM of each the forward and reverse primers. The amplification protocol consisted of 95°C for 10 min, followed by 40 cycles at 95°C for 15 s and 60°C for 60 s.

### Statistical analysis

Mann-Whitney tests were used to detect significant differences between the experimental groups. Differences were considered significant at *P* < 0.05. StatView statistical software (SAS Institute) was used for analyses. Error bars in all graphs represent S.E.M.

## Acknowledgements

We thank members of the Y.K. lab for comments on the manuscript.

## Author contributions

Conceptualization: I.H., K.K.; Methodology: H.I., W.F. K.K.; Validation: H.I., K.K.; Formal analysis: I.H.; Investigation: H.I., W.F. K.K.; Resources: I.H., W.F.; Writing - original draft: I.H., K.K.; Writing - review & editing: W. F., Y.K., K.T.K.; Visualization: I.H., K.K.; Supervision: Y.K., K.T.K.; Project administration: K.K.; Funding acquisition: K.K.

